# Spontaneous confinement of mRNA at RNP granule interfaces

**DOI:** 10.1101/2022.11.28.518040

**Authors:** Rebecca T. Perelman, Andreas Schmidt, Umar Khan, Nils G. Walter

## Abstract

Cellular membraneless organelles termed ribonucleoprotein (RNP) granules often are enriched in messenger RNA (mRNA) molecules relative to the surrounding cytoplasm. Yet, the spatial localization and diffusion of mRNAs in close proximity to phase separated RNP granules is not well understood. In this study, we performed single molecule fluorescence imaging experiments of mRNAs in live cells in the presence of two types of RNP granules, stress granules (SG) and processing bodies (PB), which are distinct in their molecular composition and function. We developed a new colocalization imaging algorithm that was employed to determine the accurate positions of individual mRNAs relative to the granule’s boundaries. We found that mRNA is often localized at granule boundaries, an observation consistent with recently published data^1,2^. We suggest that mRNA molecules become spontaneously confined at the RNP granule boundary similar to the adsorption of polymer molecules at liquid-liquid interfaces, which is observed in various technological and biological processes.

## Main text

Two U2-OS cell lines were used, stably transfected with GFP-G3BP1 and GFP-Dcp1a (termed UDG cells) to label SGs and PBs, respectively^3,4^. Cells were treated with 500 μM sodium arsenate to induce oxidative stress and form SGs^3,4^. We enzymatically synthesized a 1,600 nt long, 5’-capped, poly-adenylated and Cy5 labeled mRNA^5^ and introduced it into both cell types via bead loading^6^. Cells were then placed onto a glass bottom dish and incubated for 45 minutes in the case of U2-OS-GFP-G3BP1 cells and for 60 minutes in the case of UGD cells. During incubation, the SGs and PBs were formed while single mRNA molecules diffused throughout the cells.

We then performed single molecule mRNA imaging and RNP granule visualization via Highly Inclined and Laminated Optical Sheet (HILO) microscopy^7^ on a Nanoimager (ONI). The simultaneous acquisition of GFP and Cy5 fluorescence emitted by the granule and mRNA labels, respectively, was performed in two coregistered channels with the emitted light split via a beamsplitter onto two halves of a high-sensitivity camera (Hamamatsu sCMOS Orca flash 4 V3)^8^. The resulting hyperstacks of images contained mRNA images in one channel and the granule images in the other. Cells were kept at 37°C using an onstage incubator. The detectors’ pixel size was 134 nm and the microscope image sequences were collected every 100 ms for 20 seconds (200 images in total). Each imaging experiment was replicated once.

Representative images from a time series of a U2-OS-GFP-G3BP1 cell with SGs overlaid on the channel with mRNAs is presented in Fig. 1. The bottom panels of Fig. 1 show a continuous 20 s long image time series of the mRNA, which appears to be confined at the boundary of the SG. The mRNA follows the SG movements, demonstrating that the mRNA is in fact positioned on the granule boundary and not outside of the granule. A similar behavior can be observed in UGD cells where the mRNAs are located in the proximity of PBs. A representative image time series of a UGD cell with PBs and mRNAs is shown in Fig. 2. Again, the mRNA appears to be localized at the boundary of the PB for the duration of the experiment.

**Fig 1.**
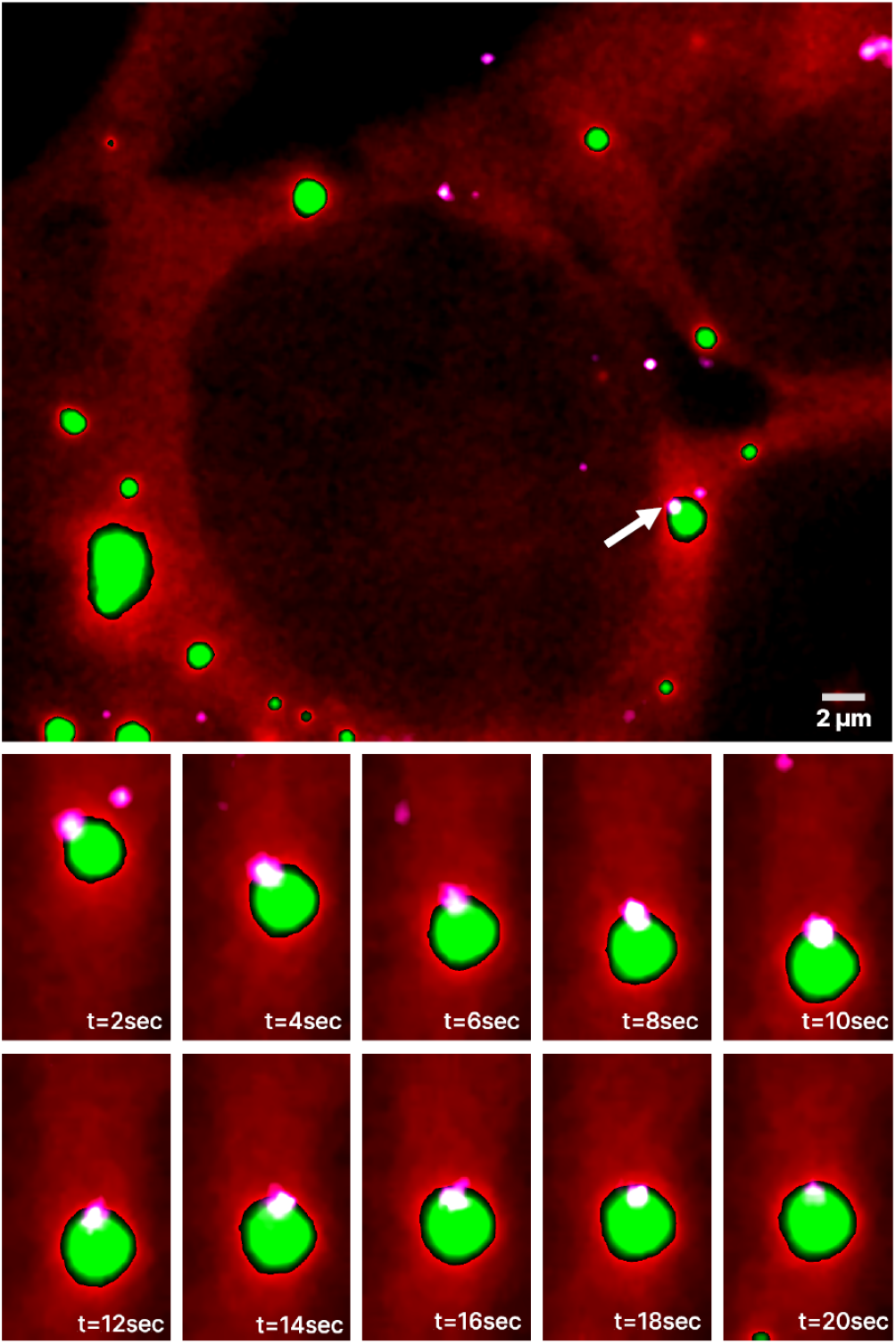
Time series of mRNAs in live U2-OS cells containing GFP labeled SGs. Representative HILO microscopy image of a U2-OS cell with GFP labeled SGs overlaid with the channel visualizing mRNAs. The two-color red-green channel visualizes SGs and U2-OS cell, green for SGs and red for U2-OS cell. The white channel visualizes mRNAs. Bottom panels show a 20s long time series of the magnified area containing mRNA, marked with a white arrow on the top panel, that appears to be confined to the boundary of the SG. The mRNA follows all SG movements.

**Fig 2.**
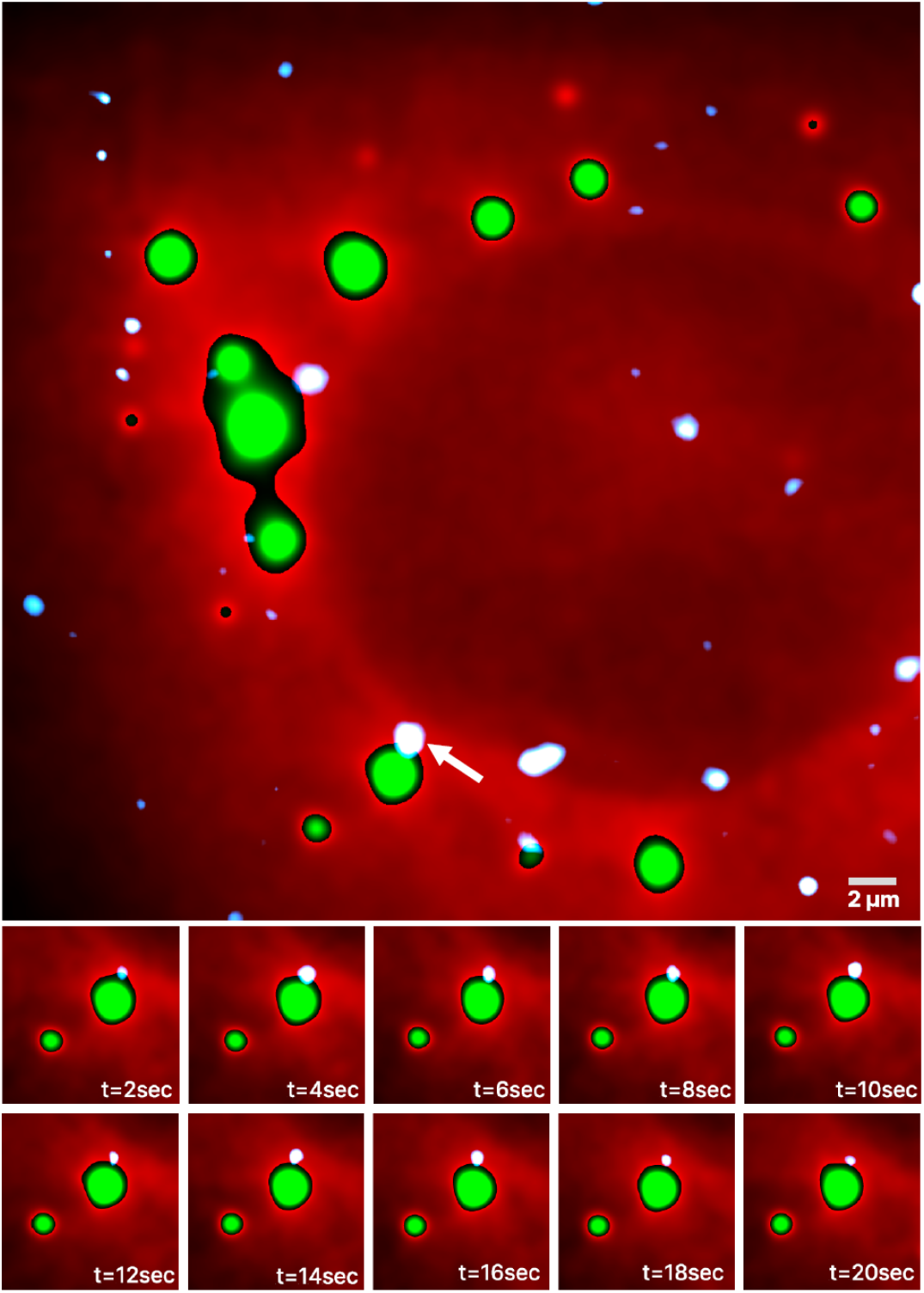
Time series of mRNAs in live UGD cells containing GFP labeled PBs. Representative HILO microscopy image of a UGD cell with GFP labeled PBs overlaid with the channel visualizing mRNAs. The two-color red-green channel visualizes PBs and UGD cell, green for PBs and red for UGD cell. The white channel visualizes mRNAs. Bottom panels show a 20s long time series of the magnified area containing mRNA, marked with a white arrow, that appears to be confined to the boundary of the PB.

Due to photobleaching, the duration of the microscopy image sequences was limited to 20 seconds and even then, the intensity of fluorescence dropped noticeably over the duration of the experiments. To address this problem, we developed an algorithm that performs photobleaching correction using a histogram matching method^9,10^. The algorithm, called image sequence colocalization with histogram matching particle detection (ISC-HMPD), was implemented as an ImageJ^11^ plugin coded in Python. In addition to photobleaching correction, ISC-HMPD reduced noise in the images with progressive switching median filters and performed mRNA molecule and RNP granule detection using the Laplacian of Gaussian (LoG) particle detection method^12^. The granules were treated as extended objects and the area of the granules was obtained from the images. As the sizes of mRNA molecules are significantly smaller than the 134 nm pixel size of the camera detector, they were treated as points. Nevertheless, due to diffraction, the mRNA images were blurred to a diffraction limited spot. To overcome the diffraction blurring, we used the fact that the signal in the mRNA single molecule detection channel comes mostly from individual, nonoverlapping mRNAs. Therefore, the coordinates of the center of the blur provide the coordinates of the mRNA with interpolation allowing for sub-diffraction accuracy. An mRNA was considered colocalized with a granule if the mRNA coordinates were within the area of the RNP granule determined with the LoG method and if the mRNA was located within ±134 nm from the granule boundary. That is, within the diffraction limit of the microscopy system, the mRNA was considered both colocalized and detected at the boundary. ISC-HMPD also computed and stored the distances between all mRNAs located within two radii from the granules and the granules’ boundaries.

In addition to the quantitative description of the mRNAs and RNP granules, ISC-HMPD allowed for the rapid processing of a large number of microscopy stacks where manual processing would be rather difficult and would risk introducing bias.

A typical output of the ISC-HMPD plugin is presented in Fig. 3 where two live U2-OS cells containing SGs (Fig. 3a) and two UGD cells containing PBs (Fig. 3b) are visualized. The algorithm determined the locations of the boundaries of the granules based on the predetermined intensity threshold. The boundaries are not shown in order to improve the visibility of the images and because the locations of the boundaries are quite obvious due to the sharp drop in intensity at the edges of the granules. Out of 14 mRNAs colocalized with the RNP granules, 12 were within ±134 nm, in other words within one diffraction limit from the boundary.

**Fig 3.**
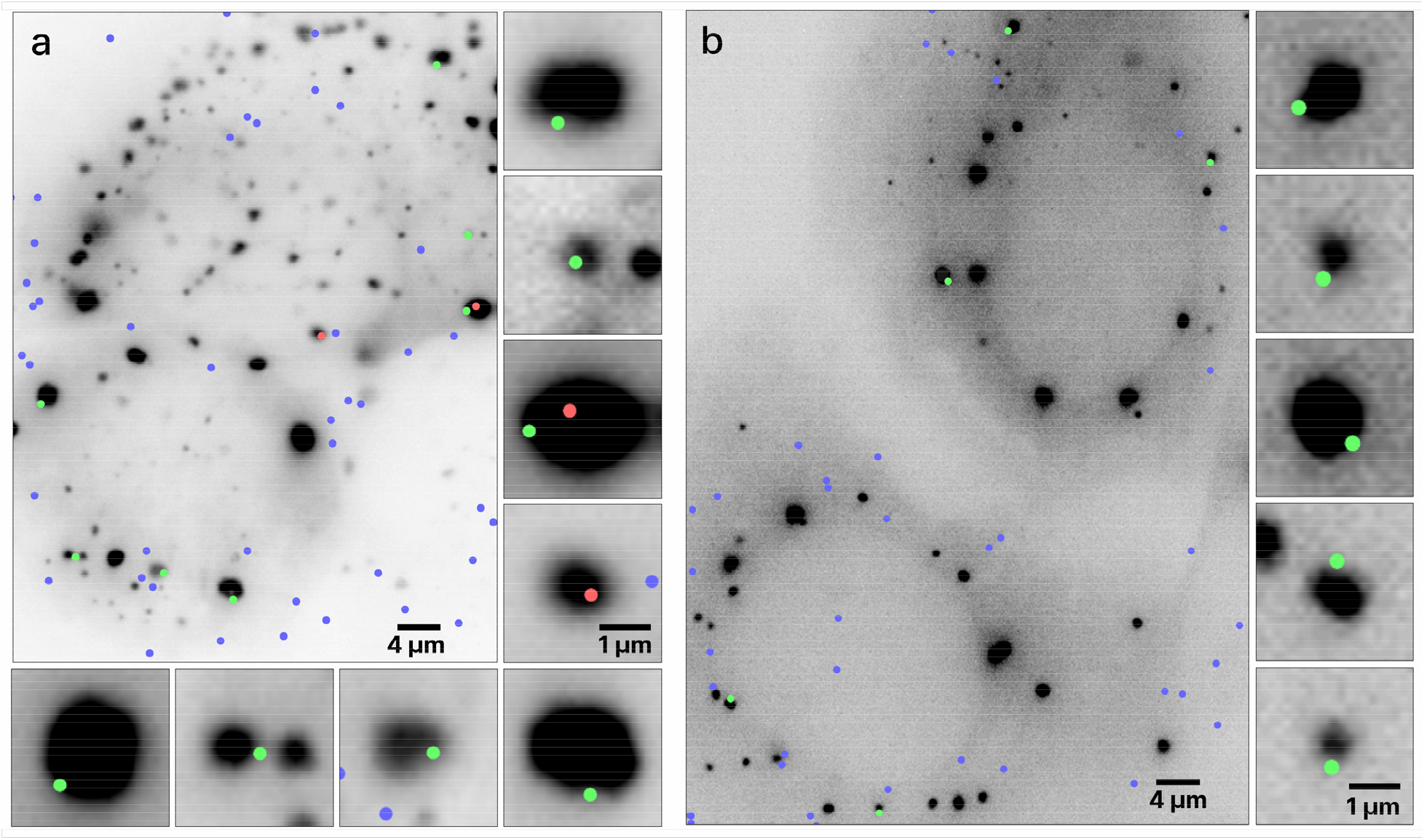
Colocalization of mRNAs and RNP granules. Two live U2-OS cells containing SGs (a) and two live UGD cells containing PBs (b) with granules imaged as dark structures. The dots represent locations of the mRNA centroids interpolated with sub-diffraction accuracy. The green and red dots represent mRNAs colocalized with the RNP granules, with the green dots specifically representing mRNAs whose centroid coordinates are within ±134 nm from the boundary. The blue dots represent the mRNAs not colocalized with the RNP granules. All mRNAs colocalized with PBs and SGs are shown on the right and bottom sides of the respective figures, approximately directly across from their location in the cell.

The algorithm determined the mRNA centroid coordinates from the single molecule HILO microscopy images and marked the locations on the cell images in color. The size of the dots was chosen based on visibility considerations and does not represent the size of the mRNA molecule, which was assumed to be much smaller than the pixel size. The center of the dot, coinciding with the maximum fluorescence intensity of the mRNA label, is the centroid of the mRNA. The mRNA, whose centroid coordinates were inside or on the boundary of the RNP granules are shown as red and green dots, respectively, and the remainder of the mRNAs are shown as blue dots.

All mRNAs colocalized with SGs in Fig. 3a are shown the right side of the figure. It appears that in this particular image, out of the 9 colocalized mRNAs, 7 were located within the diffraction limited distance from the boundary of the granule, and therefore, on the boundary and 2 inside the SG. A similar observation holds for PBs in Fig. 3b, where all 5 colocalized mRNAs are located on the boundary.

The images are quite representative of all of our data, in terms of mRNAs predominantly being found at the granule boundaries in both UGD and U2-OS cells. Overall, we determined that out of approximately 10,000 mRNAs imaged in the UGD and U2-OS cells, around 80% were located on the boundaries of the PBs and SGs. Furthermore, as the microscopy system has a final depth of view, some of the mRNA molecules that are actually located on the boundary of the granule may appear to be localized to the inside of the granule in the microscopy images. Therefore, these percentages represent a lower boundary of the fraction of mRNAs that are localized to the granule boundaries.

We also determined the average spatial density distribution of the mRNAs relative to the boundary of the nearest RNP granule (Fig. 4) with the distance measured to the closest point of the granule. The negative distances represent the locations outside the granule and positive distances represent the inside of the granule. In both PBs and SGs, we clearly observed a peak of mRNA density distribution at the boundaries of the granules.

**Fig 4.**
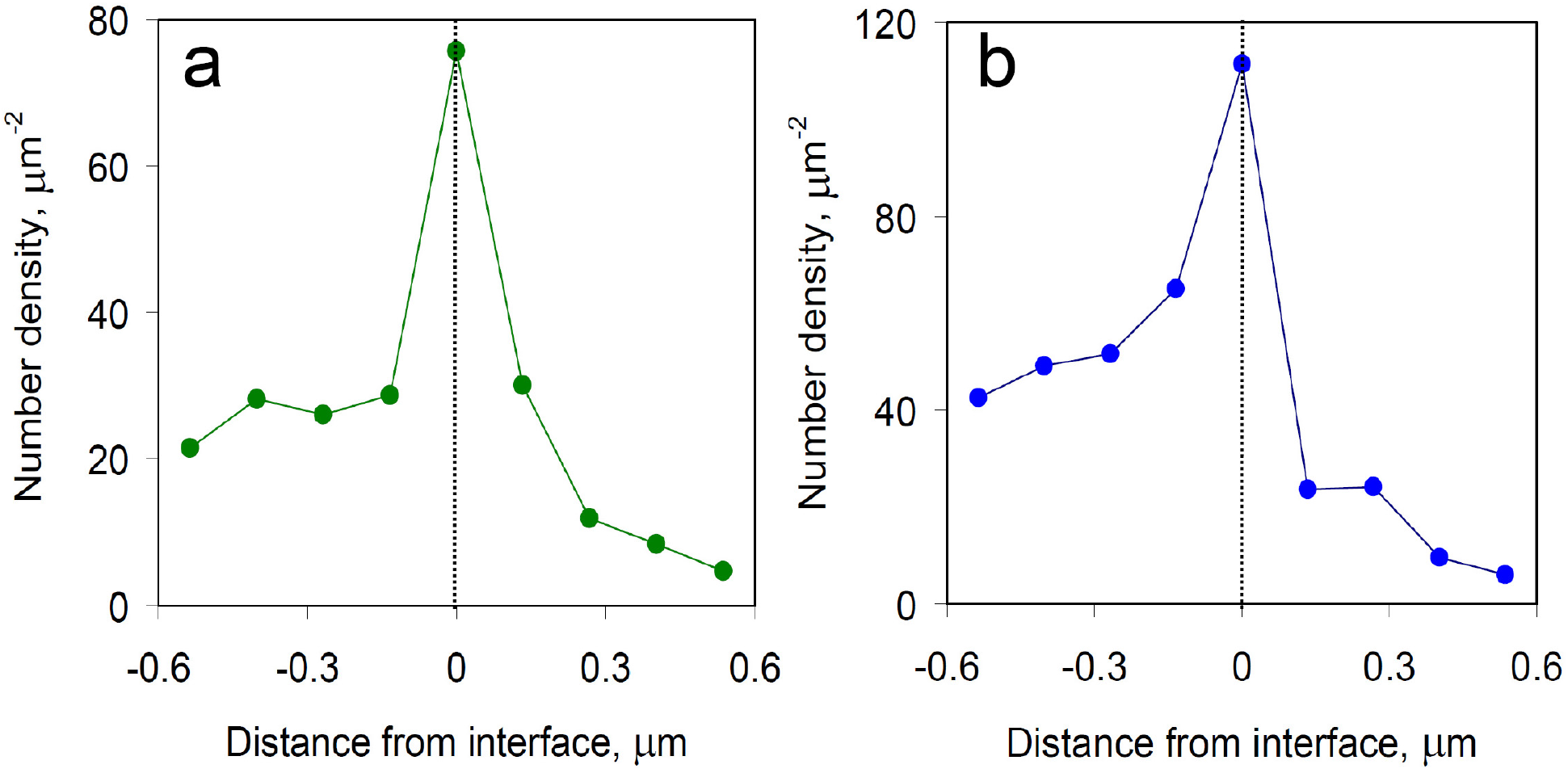
Spatial density distribution of the mRNAs relative to the boundaries of the nearest RNP granules. Average spatial density distribution of the mRNAs relative to the boundaries of the nearest SG (a) and PB (b). Negative distances represent the locations outside of the granule and positive distances represent the inside, the dashed line marks the location of the boundary.

These observations are consistent with data from recent studies that also observed that mRNAs are predominantly found at the boundaries of the biomolecular condensates^1,2,4^. For example, in a recent study employing imaging in the visible wavelength range with confocal fluorescence microscopy of mRNAs in SGs, over two-thirds of the mRNAs were located within the optical resolution of the microscope, at the granules’ boundaries^2^.

Apart from the imaging experiments described here, which determined that mRNAs are predominantly localized at the RNP granule boundaries, there are other indirect observations that support these observations. Recently, it was observed that mRNAs imaged in close proximity of the RNP granules can be described with at least two characteristic times^1^: a fast time on the order of seconds^13-16^ and a slow time where mRNA becomes “locked” to the RNP granule for tens of minutes and even hours^1^. The consensus is that one of these times is likely related to the diffusion of mRNA within the RNP granule. However, the nature of the other characteristic time is unclear. One potential explanation for the existence of several characteristic times may be related to the known inhomogeneity of RNP granules^17^. However, such an inhomogeneity would likely result not in two characteristic times, but in multiple times associated with the mRNA traveling across dense cores of various sizes and also the various distances between these cores^1^. On top of that, these times would likely differ not just among different types of biomolecular condensates but even in the same type of condensate. Therefore, it seems more likely that one of these characteristic times is associated with the spontaneous confinement of the mRNA at the RNP granule interface.

To determine whether this confinement is the fast or slow time, we estimated the characteristic time, *τ*_*d*_, of mRNA diffusing across an RNP granule of size *d*. The diffusion of mRNA can be described using Brownian motion in a highly viscous liquid. From Einstein’s well known expression for the mean squared displacement in three dimensions: ⟨**r**^2^⟩ = 6*Dτ_d_*, where *D* is the diffusion coefficient of mRNA in an RNP granule, and assuming that on average 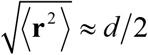, using the standard Einstein - Stokes equation for the diffusion coefficient, we get *τ* ≈ *π d* ^2^*η R/*4 *k*_*B*_*T*, where *R*_*g*_ is the radius of gyration of mRNA, *T* is the temperature, and *k*_*B*_ is the Boltzmann constant. The majority of the parameters in this formula vary within a rather narrow range, making the viscosity *η* of the gel-like medium inside the RNP granule the single most important parameter that can affect the characteristic time of the diffusion of mRNA within an RNP granule. Experimental measurements in several types of biomolecular condensates, including P granules^18^, TDP-43 RNP granules^19^, PGL-3 condensates^20^, and condensates formed in LAF-1^18,21^ yielded viscosities in the range of 0.1 to 34 Pa·s. Even for the highest viscosity in that range at room temperature, for a typical mRNA with a radius of gyration of 10 nm and a relatively large, one micrometer in size granule, the characteristic diffusion time would be approximately two minutes. This is an order of magnitude shorter than the 20 to 60 minutes previously observed experimentally^1^. Therefore, in order to associate the slow time with the diffusion of mRNA within the RNP granule, the viscosity in the RNP granule should be at least 400 Pa·s. This value is at least an order of magnitude larger than those reported for various types of biomolecular condensates, including RNP granules. Therefore, it is reasonable to assume that for an mRNA localized to an RNP granule to have a second characteristic time while only having two characteristic times, there should be another mechanism responsible which is not directly related to diffusion. It appears that it is more likely that the fast time is associated with the diffusion of mRNA within the RNP granule. Therefore, the slow time should then be associated with the spontaneous confinement of mRNA at the RNP granule boundary due to an interaction with the liquid-liquid interface.

Messenger RNA is a polyanionic molecule. Polymers can experience confinement or adsorption at the interface of two liquids. This phenomenon has been extensively studied experimentally, theoretically, and computationally^22-24^. The desorption time is a parameter which characterizes the degree of polymeric molecule confinement and can be measured experimentally. The desorption time of a single polymer molecule from liquid-liquid interfaces has been recently modeled computationally^25^. A possible explanation for polymer adsorption could involve the molecular-level amphiphilicity of the polymer. However, a more general explanation could be related to the fact that the presence of a polymer molecule at the liquid-liquid interface can locally lower the interfacial free energy, trapping the polymer molecule at the interface.

In conclusion, we observed the confinement of mRNAs at the interfaces of two types of RNP granules, SGs and PBs. We believe this effect could be common for all types of biomolecular condensates. We suggest that the spontaneous confinement of mRNA at the RNP granule boundaries is similar in nature to the adsorption of polymer molecules observed in various technological and biological processes. Additional studies are necessary to confirm the commonality of mRNA confinement at the interfaces of other types of biomolecular condensates and to establish which interaction mechanism is primarily responsible for the confinement.

## Methods

### Cell line cultivation, handling and imaging

U2-OS (HTB-96, ATCC), stably expressing GFP-G3BP1, were maintained in DMEM and U2-OS-GFP-DCP1A (UGD) cells were maintained in McCoy’s 5A medium (GIBCO). Both cells were supplemented with 10% (v/v) fetal bovine serum (GIBCO) and 1% (w/v) Penicillin-Streptomycin (GIBCO) at 37°C under 5% CO2. UGD cells were kept under positive selection with 100 μg/mL G418^26^. U2-OS cells stably expressing GFP-G3BP1 were a gift from the Moon lab^1^. Oxidative stress was induced in U2-OS cells by treating them with 0.5 mM sodium arsenite (NaAsO2, SA) for 60 min to form SGs.

Messenger RNAs were loaded into cells via bead loading^27^. Briefly, medium was removed, and cells were washed 2 times with 1 ml 1x PBS buffer, 5 µl of RNA solution in 1x PBS buffer (200 ng) was added to the center of the glass dish and then glass beads were also added. Afterwards, the glass dish containing glass beads was tapped 10 times against the bench and the culture medium was then added back in and incubated at 37 °C for 60 min (SG experiments) and 75 min (PB experiments) before imaging. For live cell imaging, cells were imaged in phenol-red free Liebovitz’s L-15 medium containing 1% FBS using a Nanoimager S from the Oxford Nanoimaging Limited (ONI) system in highly inclined laminated optical sheet (HILO) illumination mode^7^. Alexa Fluor 647 dye was detected using a 640 nm red laser and GFP using a 473 nm laser. The integration time was 100 ms.

### Generation of fluorophore labeled mRNA

The synthesis included three steps^5^. First, an enzymatic RNA synthesis was performed in a run-off in vitro transcription using T7 RNA Polymerase (ThermoFisher Scientific) as described by the manufacturer. DNA transcription templates were generated from a corresponding plasmid by PCR. Five % (v/v) of the unpurified PCR product was used as a template in the in vitro transcription. The reaction was incubated at 37°C for 90 minutes. Following ethanol precipitation, RNA was resuspended in double distilled H_2_O. Secondly, the 3’-end azide functionalization in presence of 2’-azido-2’-deoxyadenosine-5’triphosphate (ATP-azide, Trilink Biotechnologies) using yeast poly(A) polymerase (yPAP, ThermoFisher Scientific) and capped using ScriptCap m^7^G Capping System (CellScript). The reaction mixture contained 1.4 µM of RNA transcript, 600 µM SAM, 1mM GTP, 700 µM ATP-azide, 10 U capping enzyme, 2400 U of yPAP and 1x Script capping buffer. Following an incubation of 60 min at 37°C, the reaction mixture was ethanol precipitated and resuspended in double distilled H_2_O. Lastly, recovered RNA was polyadenylated in the presence of ATP using 2400 U yeast poly(A) polymerase (yPAP, ThermoFisher Scientific) in 1x Poly(A) polymerase reaction buffer for 30 min at 37°C and then, in the same mix, labeled using 150 molar excess of Click-IT Alexa Fluor 647 sDIBO Alkyne (ThermoFisher Scientific) at room temperature for 60 min.

### Image Sequence Colocalization

The acquired image sequence consisted of the HILO microscopy-based hyperstacks of images with one of the channels containing mRNA images and the other containing the granule images. The images were processed with image sequence colocalization with the histogram matching particle detection (ISC-HMPD) algorithm which we developed and implemented in Python. The processing pipeline included the photobleaching correction with histogram matching, noise removal with progressive switching median filters, Laplacian of Gaussian-based particle detection, and colocalization of all detected and isolated granules and mRNA molecules.

The channel containing the single molecule mRNA images exhibited a significant decay in signal strength over time due to photobleaching. To compensate for this continuous decay, we employed a histogram matching transformation^9,10^. Histogram matching involves the manipulation of pixels of an unprocessed image or sequence of images to match them with a reference image.

This is done by finding the cumulative histogram of both the reference and original image. Then the cumulative histogram of the original unprocessed image(s) is adjusted to match the unprocessed image(s) with the cumulative distribution function of a given reference image. Figure 5 provides an example of the histogram matching transformation performed on one of the frames from our image sequence. The correction for photobleaching can be seen when an image at 10s lacks contrast; however, the histogram matching algorithm corrects the contrast by using the reference image at 0s which results in a high-contrast image at 10s.

**Fig 5.**
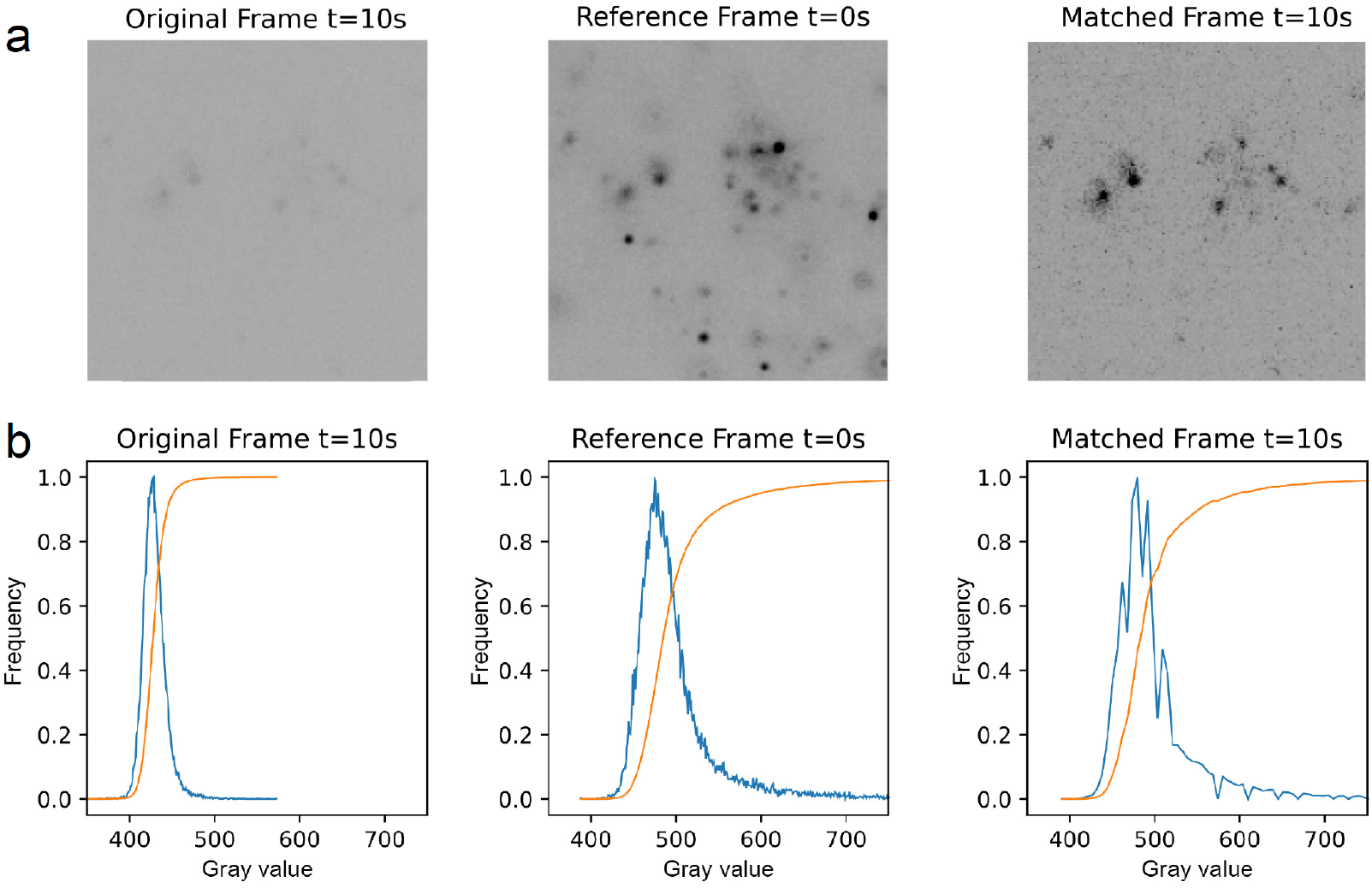
Example illustrating histogram matching gray value adjustment of an arbitrary frame in an image sequence using the first frame as a reference. (a) Original frame at 10s lacks contrast, while revised matched frame at 10s shows improved contrast when its histogram is matched with the reference frame at 0s. (b) Plots showing histograms (in blue) of respective frames above and cumulative distribution functions (in orange).

In order to detect mRNA and granules and to also obtain the granules’ area in the image sequence dataset, we implemented the scale-normalized Laplacian of Gaussian (LoG) detection algorithm, which is well-suited for our application due to its strong response to particles residing in the inhomogeneous background^27^. The pixel size of the image was 134 nm. An mRNA was determined to be colocalized with a granule if the coordinates of the mRNA were within the area of the granule determined with the LoG algorithm. If the mRNA was within ±134 nm from the boundary, or in other words within the diffraction limit of the microscopy system from the boundary, the algorithm characterized it as located at the boundary and presented it as a green dot with its coordinates located a the centroid of the mRNA blur. The rest of the colocalized mRNAs were presented as red dots and non-colocalized mRNAs were presented as blue dots.

## Statistical Details

Statistical details such as the number of mRNAs analyzed are indicated in the text and figure legends.

## Acknowledgments

This work was supported by NIH R35 grant GM131922 to N.G.W. and the Deutsche Forschungsgemeinschaft (DFG, German Research Foundation) Project-ID 449562930 to A.S. We thank L.T. Perelman for input on the manuscript.

## Author contributions

R.T.P. conceived the method, performed laboratory experiments, developed a physical model, programmed the algorithm, performed the data and statistical analysis, prepared the figures, and wrote the manuscript; A.S. performed laboratory experiments; U.K. programmed the algorithm, and prepared figures; N.G.W. initiated and supervised the project, evaluated the method, and edited the manuscript.

## Additional information

Correspondence and requests for materials should be addressed to N.G.W.

